# Network Mechanisms Underlying the Role of Oscillations in Cognitive Tasks

**DOI:** 10.1101/271973

**Authors:** Helmut Schmidt, Daniele Avitabile, Ernest Montbrió, Alex Roxin

**Affiliations:** Centre de Recerca Matemàtica, Campus de Bellaterra Edifici C, Bellaterra, Barcelona, Spain; Barcelona Graduate School of Mathematics, Campus de Bellaterra Edifici C, Bellaterra, Barcelona, Spain; School of Mathematical Sciences, University of Nottingham, University Park, Nottingham, United Kingdom; Center for Brain and Cognition, Department of Information and Communication Technologies, Universitat Pompeu Fabra, Barcelona, Spain

## Abstract

Oscillatory activity robustly correlates with task demands during many cognitive tasks. However, not only are the network mechanisms underlying the generation of these rhythms poorly understood, but it is also still unknown to what extent they may play a functional role, as opposed to being a mere epiphenomenon. Here we study the mechanisms underlying the influence of oscillatory drive on network dynamics related to cognitive processing in simple working memory (WM), and memory recall tasks. Specifically, we investigate how the frequency of oscillatory input interacts with the intrinsic dynamics in networks of recurrently coupled spiking neurons to cause changes of state: the neuronal correlates of the corresponding cognitive process. We find that slow oscillations, in the delta and theta band, are effective in activating network states associated with memory recall by virtue of the hysteresis in sweeping through a saddle-node bifurcation. On the other hand, faster oscillations, in the beta range, can serve to clear memory states by resonantly driving transient bouts of spike synchrony which destabilize the activity. We leverage a recently derived set of exact mean-field equations for networks of quadratic integrate-and-fire neurons to systematically study the bifurcation structure in the periodically forced spiking network. Interestingly, we find that the oscillatory signals which are most effective in allowing flexible switching between network states are not smooth, pure sinusoids, but rather burst-like, with a sharp onset. We show that such periodic bursts themselves readily arise spontaneously in networks of excitatory and inhibitory neurons, and that the burst frequency can be tuned via changes in tonic drive. Finally, we show that oscillations in the gamma range can actually stabilize WM states which otherwise would not persist.

**Author Summary:** Oscillations are ubiquitous in the brain and often correlate with distinct cognitive tasks. Nonetheless their role in shaping network dynamics, and hence in driving behavior during such tasks is poorly understood. Here we provide a comprehensive study of the effect of periodic drive on neuronal networks exhibiting multistability, which has been invoked as a possible circuit mechanism underlying the storage of memory states. We find that oscillatory drive in low frequency bands leads to robust switching between stored patterns in a Hopfield-like model, while oscillations in the beta band suppress sustained activity altogether. Furthermore, inputs in the gamma band can lead to the creation of working-memory states, which otherwise do not exist in the absence of oscillatory drive.

## Introduction

Oscillations are ubiquitous in neuronal systems and span temporal scales over several orders of magnitude [1]. Some prominent rhythms, such as occipital alpha waves during eye-closure [2] or slow-oscillations during non-REM sleep [3] are indicative of a particular behavioral state. Other rhythms have been specifically shown to correlate with memory demands during working memory tasks, including theta (4 - 8Hz) [4–7], alpha/beta (8 - 30Hz) [8–10] and gamma (20 - 100Hz) [11–13]. Nonetheless, neither the physiological origin nor the functional role of such oscillations are well understood.

Here we study how oscillatory signals in distinct frequency bands can serve to robustly and flexibly switch between different dynamical states in cortical circuit models of working memory and memory storage and recall. In doing so we characterize the dynamical mechanisms responsible for some of the computational findings in an earlier study [14]; we go beyond that work to include new results on oscillatory control of network states. Specifically, we consider the response of multistable networks of recurrently coupled spiking neurons to external oscillatory drive. We make use of recent theoretical advances in mean-field theory to reduce the spiking networks to a low-dimensional macroscopic description in terms of mean firing rate and membrane potential, which is exact in the limit of large networks [15]. This allows us to perform a systematic and detailed exploration of network states analytically or with numerical bifurcation analysis, which informs us about suitable parameter sets for numerical simulations. The latter serve to give representative examples of the dynamical phenomena investigated here. As a result, we can completely characterize the dynamics of the forced system.

Specifically, we consider networks which exhibit multistability in the absence of forcing. Such attracting network states have been proposed as the neural correlate of memory recall [16,17], and as a possible mechanism for sustaining neuronal activity during working memory tasks [18–20]. We find that an external oscillatory drive interacts with such multistable networks in highly nontrivial ways. Low-frequency oscillations are effective in switching on states of elevated activity in simple bistable networks, while in higher dimensional multistable networks they allow for robust switching between stored memory states. Higher frequencies, in the beta range, destabilize WM states through a resonant interaction which recruits spike synchrony. Such oscillatory signals can therefore be used to clear memory buffers. Finally, when networks operate outside the region of multistability, e.g. due to reduced excitability, an oscillatory signal in the gamma range can be used to recover robust memory recall.

## Results

### Oscillatory drive can selectively turn on or off WM states

Networks of recurrently coupled excitatory neurons can exhibit bistability given sufficiently strong synaptic weights. Such networks act as binary switches: a transient input can cause a transition from a baseline state to a state of elevated activity, or vice-versa. We asked to what extent an oscillatory signal alone could also drive transitions between states in such a network. In particular we were interested in knowing if the directionality of the transition, and hence the final state of the system, could be controlled via the frequency of the oscillatory drive.

To investigate this we simulated a network of recurrently coupled excitatory quadratic integrate-and-fire neurons, see Materials and Methods for details. Fig 1 shows an illustration of the network dynamics as a function of the stimulus frequency and initial state of the network. In particular, at low frequencies, the oscillations push the system from the state of low activity into the state of high activity, which persists under such forcing, see Fig 1A. As the frequency is increased past a critical value, it is no longer effective in driving a transition, and the network remains in its initial state, see Fig 1B. A further increase then shows the opposite effect: The state of high activity becomes unstable under the forcing, whereas the state of low activity persists, Fig 1C. At large enough frequencies we then observe again that no transitions occur and the initial network state persists, Fig 1D.

**Fig 1.**
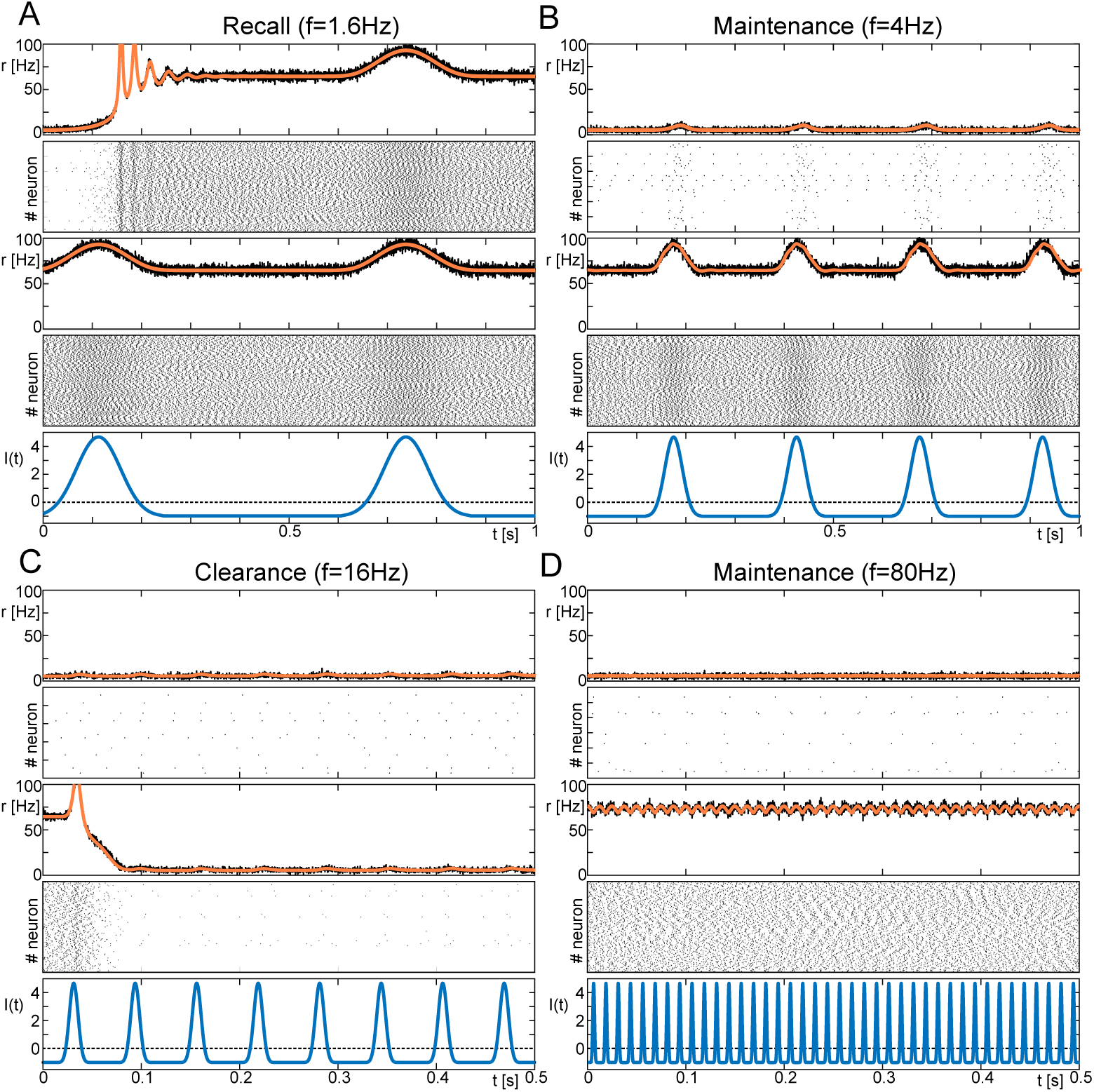
Frequency response of a network of QIF neurons. Here we show the response of a network of 10^4^ all-to-all coupled QIF neurons with distributed input currents to periodic forcing. The model parameters are chosen such that the network is bistable, see also Fig 2A. Each panel shows the network-averaged firing rate and raster plot of the response for an initial condition in the low-activity state (top, *r* ≈6Hz) and high-activity state (bottom, *r*≈73Hz). **A** At low enough frequencies, the system is pushed from the low-to the high-activity state. **B** At slightly higher frequencies, both states persist under the forcing. **C** Driven with frequencies from an intermediate range of frequencies, the state with high firing activity destabilizes in favor of the state with low firing activity. **D** At high frequencies, both states persist under the forcing. Parameters: 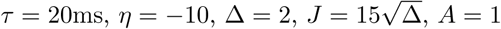.

The results from Fig 1 show that the frequency of an external oscillatory drive can be used to selectively stabilize a given network state. For the parameter values used here, oscillation frequencies in the delta range result in a WM state while frequencies in the beta range force the system to the “ground” state, essentially clearing the WM state, a result seen also in [14]. Oscillations outside these ranges are ineffective in driving transitions. We seek to understand the mechanisms underlying these transitions, and additionally to determine to what extent the precise frequency ranges are influenced by the network parameters. To do this we will take advantage of recent work in which the authors derived a set of simple equations for the mean firing rate and mean membrane potential in a network of recurrently coupled quadratic integrate-and-fire (QIF) neurons [15]. In the large-system limit these equations are exact. The exact correspondence between the low-dimensional mean-field equations and the original network allows us to use standard dynamical systems techniques to fully characterize the range of dynamical states in the network.

### Model equations and network analysis

The dynamics in networks of recurrently coupled QIF neurons can be described exactly under the assumptions of all-to-all coupling and quenched neuronal variability, i.e. static distributions in cellular or network properties. For the case of a single network of excitatory cells in which the input currents to individual neurons are distributed, the resulting mean-field equations are [15]:

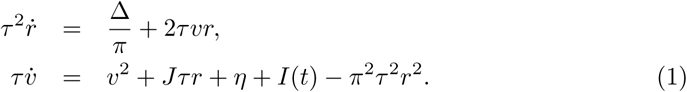

Here, *r* is the network average of the firing rate and *v* is the network average of the membrane potential, *J* is the strength of synaptic weights, *η* is the mean of the static distribution of inputs, while Δ is the width of the input distribution. External, time-variant forcing is represented here by *I*(*t*). The time constant *τ* is the membrane time constant of the individual neurons and is set to 20ms throughout.

This macroscopic model permits a straightforward investigation of the stationary states in the full network. For sufficiently strong synaptic coupling two stable fixed points co-exist over a range of mean external inputs, see Fig 2A (left). Linear stability analysis further reveals that the stable high-activity fixed point is a focus for sufficiently high rates, whereas the stable low-activity fixed point is a node, see Materials and Methods. The network therefore shows a damped oscillatory response to external perturbations in the high-activity state. This response reflects transient spike synchrony which decays over time due to the heterogeneity; the characteristic time scale of the desynchronization is in fact proportional to the width of the distribution of input currents Δ. This type of spike synchrony is seen ubiquitously in networks of heterogeneous and noise-drive spiking neurons [15,21,46] and is captured in Eq 1 by the interplay between the mean sub-threshold membrane potential and mean firing rate [23].

**Fig 2.**
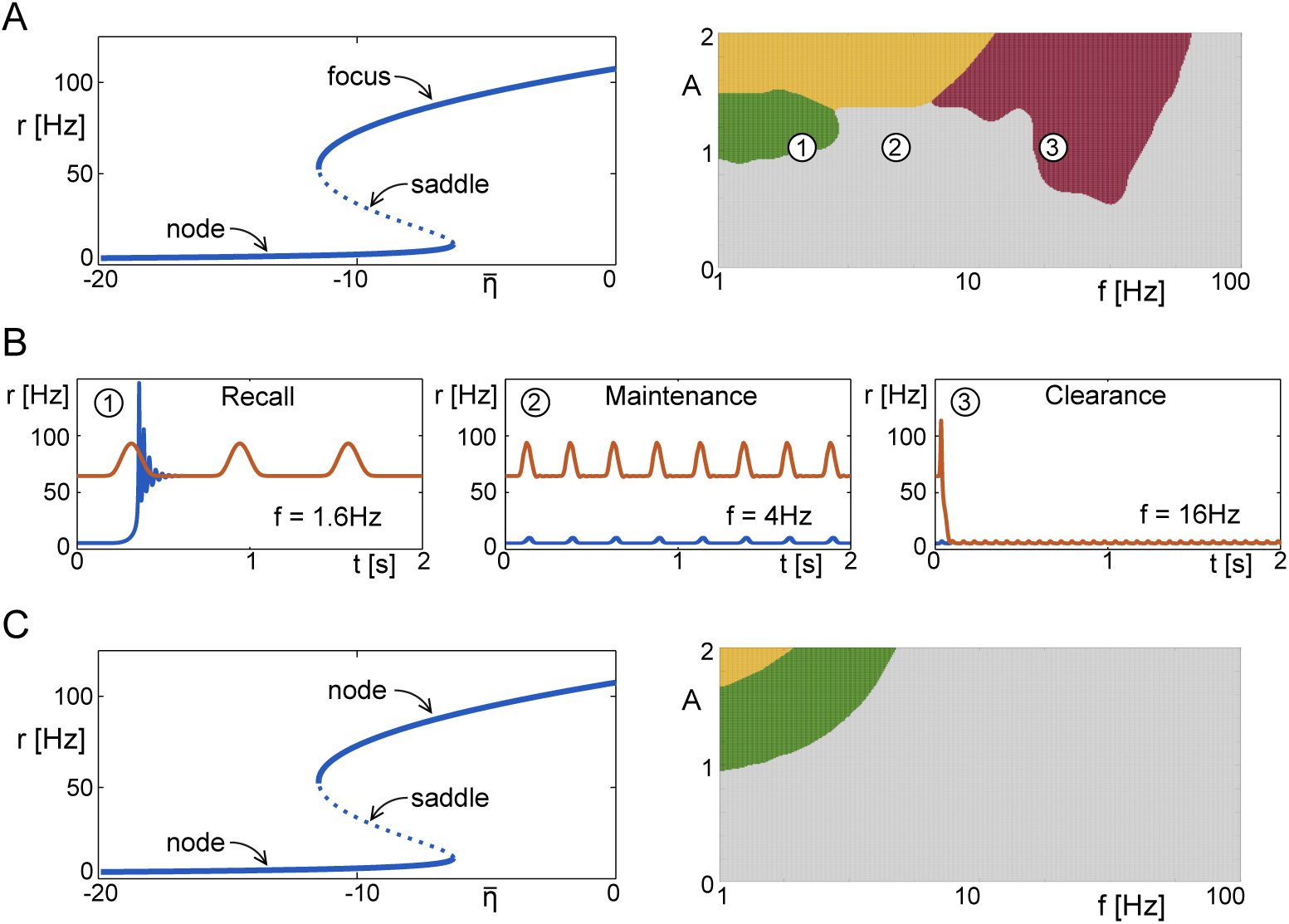
Switching behavior at the macroscopic scale. **A** Bifurcation analysis identifies a bistable regime where a stable focus and a stable node coexist. The different dynamic regimes are shown here as a function of the amplitude *A* and the frequency *f* of the forcing. Green: Recall; Grey: Maintenance; Red: Clearance. Orange: only one globally stable limit cycle exists due to the system being entrained to the forcing. **B** Example time traces from **A**, with initial conditions chosen to be the focus (red) or the node (blue). **C** The heuristic firing-rate equations Eq. 4 with equivalent fixed point structure do not show the same dynamic regimes, as the focus is reduced to a node and therefore cannot be destab<u>il</u>ised by nonlinear resonance. Parameters: 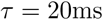 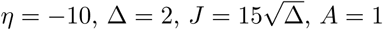.

We use this macroscopic description to systematically investigate the network response to periodic forcing with amplitude *A* and frequency *f*. Fig 2A (right) shows a phase diagram of the network dynamics as a function of these two parameters. As in Fig 1 we keep track of the final state of the network as a function of the initial state. For sufficiently slow frequencies and over a range of amplitudes the network is always driven to the high-activity state (green). This region therefore corresponds to recall of the memory state, see Fig 2B (left). For an intermediate range of frequencies a suffienctly strong forcing always drives the network to the low-activity state (red), which corresponds to clearance, Fig 2B (right). The frequency band for clearance is essentially set by the frequency of intrinsic oscillations of the high-activity state, i.e. it is a resonant effect, see Fig 3. Weak forcing and forcing at very high frequencies fail to drive any transitions, while strong forcing at low enough frequencies can enslave the network dynamics entirely (orange). For the parameter values used here recall occurs for frequencies below about 2Hz and clearance in the range between 10-30Hz.

**Fig 3.**
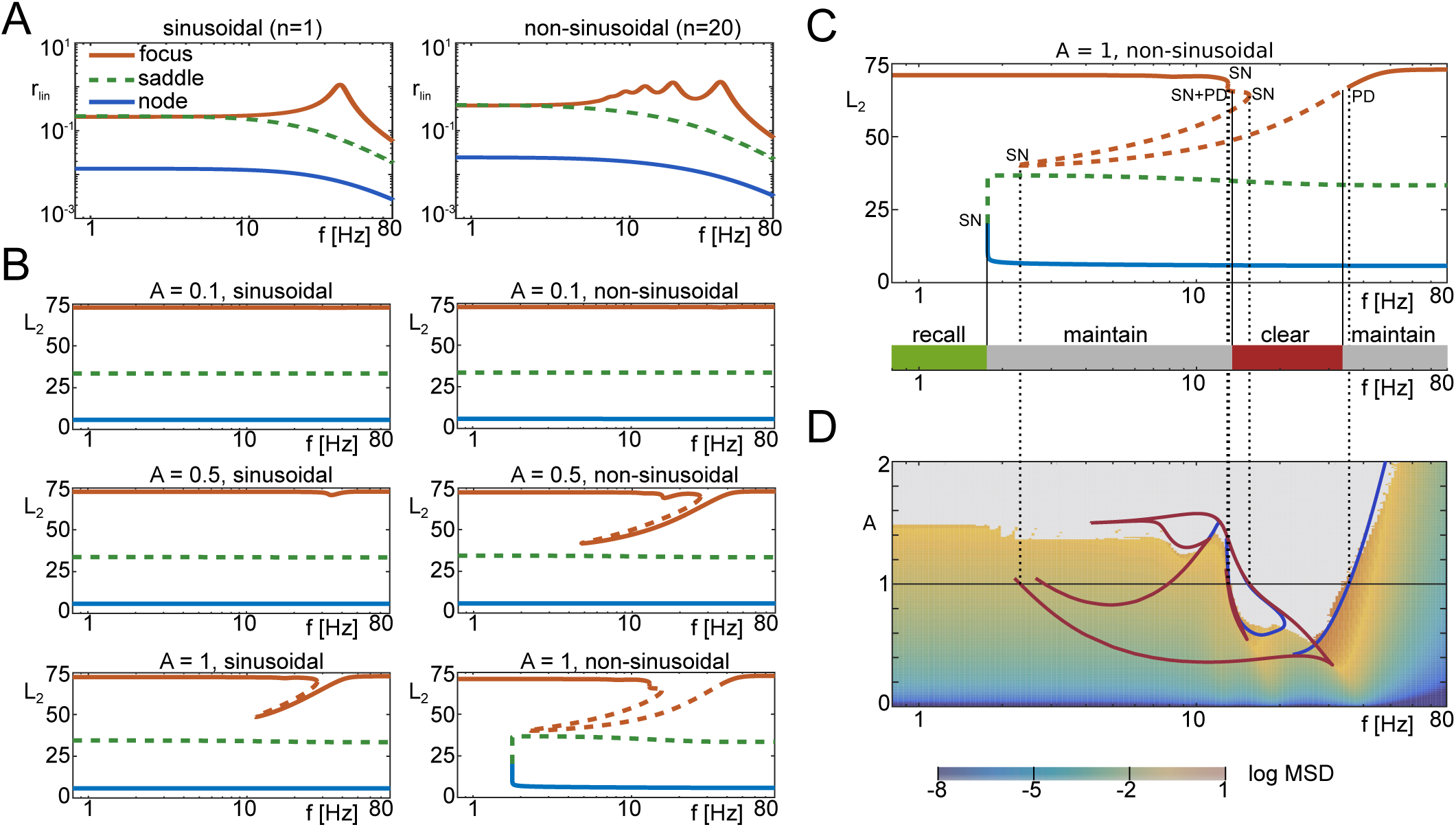
Linear and nonlinear response. **A** Linear response of focus, saddle and node to sinusoidal and non-sinusoidal inputs, with the focus showing a characteristic resonant response at approximately 40Hz. The response of the focus to non-sinusoidal input shows additional sub-harmonic resonances. **B** Nonlinear response of the fixed points by means of bifurcation analysis in the forcing frequency for different amplitudes. Non-sinusoidal forcing leads to a richer bifurcation structure. **C** Bifurcation diagram in *f* for non-sinusoidal forcing with *A* = 1, and comparison with numerical results (bottom). The bifurcation structure is governed by saddle-node bifurcations (SN) and period-doubling bifurcations (PD). Branches of period-doubled solutions are omitted here. **D** A two-parameter bifurcations analysis of the focus reveals the loci of saddle-node bifurcations (red) and period-doubling bifurcations (blue) in the (*f, A*)-plane. We compare these with the logarithmic mean squared deviation (log MSD) from the fixed point (color scale), obtained by time simulations. Grey areas indicate regions where the system leaves the basin of attraction of the focus. Parameters: 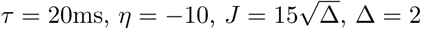.

In order to characterize the role of spike synchrony in determining the network response, we derive a reduced firing rate equation with the identical fixed-point structure as in the original, exact mean-field equations Eq 1, but without the subthreshold dynamics. Specifically, the fixed-point value of the firing rate in Eq 1 can be written as

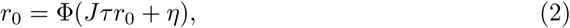

where Φ is the steady-state f-I curve, which in the case of Eqs (1) is

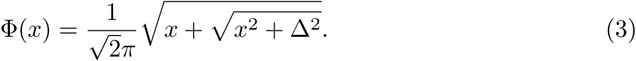

We use the steady-state f-I curve to construct a heuristic firing rate model given by

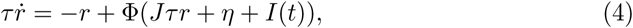

and investigate its response to periodic forcing *I*(*t*), see Materials and Methods for more details. In this case the high-activity branch of the firing rate is a node, i.e. it no longer shows damped oscillations in response to perturbations, see Fig 2C (left). Furthermore, the region of “clearance” has completely vanished in the phase diagram in Fig 2C (right), confirming that in the original network it was due to a resonance reflecting an underlying spike synchrony mechanism.

### Recall and Clearance occur due to forcing-induced bifurcations

Given the simplicity of the mean-field equations Eq 1 we can calculate the linear response of the system analytically, without the need for extensive numerical simulations. The response of the focus to weak sinusoidal inputs (linear response) already shows a clear resonance for the high-activity state (Fig 3A), where the resonant frequency is

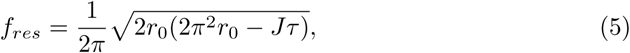

see Materials and Methods. Furthermore, additional, sub-harmonic resonance peaks occur when the forcing is sharply peaked, leading to a broadening of the resonance spectrum (Fig 3A, right); this effect is due to the presence of many sub-harmonics of the linear resonance in the forcing term itself. Conversely, the node does not show such a resonance, indicating a qualitative difference in the response of the two stable fixed points.

However, the switching behaviors seen in Fig 1 and Fig 2 and the corresponding destabilization of network states cannot be attributed to this linear resonance alone – nonlinear effects have to be taken into account. This can be seen by plotting the bifurcation diagram for the response of the network to the forcing for several values of the forcing. For relatively weak, but finite forcing, the network response consists of a limit cycle in the vicinity of the corresponding unforced fixed-point, Fig 3B (top). As the forcing amplitude is increased the amplitude of the limit cycle inherited from the high-activity state is selectively enhanced due to the resonance, eventually leading to bistability of limit cycles over a range of frequencies, see Fig 3B middle-right and bottom-left.

At large enough amplitudes for the sharply-peaked, non-sinusoidal forcing two additional bifurcations occur which are responsible for the “recall” and “clearance” behaviors respectively, see Fig 3B (bottom-right) and Fig 3C. Specifically, at low frequencies the two limit cycle solutions which arise due to the periodic forcing of the low-activity node (blue line) and saddle-point (green line) anhihilate in a saddle-node bifurcation of limit cycles. Therefore at low frequencies the only stable solution is the limit cycle in the vicinity of the high-activity focus (red line), see Fig 3C. This explains why low frequencies are effective in switching on the high-activity state, i.e. for “recall”. On the other hand, in the range of frequencies over which the network response is resonant, period-doubling bifurcations of the focus lead to a frequency band in which all responses of the focus are unstable. Therefore, the limit-cycle solution in the vicinity of the low-activity node is the only stable solution. Frequencies in this range are therefore effective in switching off the high-activity state, i.e. for “clearance”. See also Supporting Information (S1 Text). The “clearance” mechanism can be reproduced in a a simplified model, see Supporting Information (S2 Text).

### Higher-dimensional memory circuits

A single bistable network of neurons serves as a canonical illustration of a memory circuit. However, such a network can only store a single bit of information; actual memory circuits must be capable of storing more information. In terms of neuronal architecture this can be achieved by having a network which is comprised of several or many neuronal clusters [16,17,24]. We asked to what extent the frequency-selective switching behavior seen in a single bistable network could also be found in a clustered network. We look first at a simple, two-cluster network and then the more general case of a higher-dimensional multi-clustered network.

#### Two competing neuronal populations

We set up a network of two identical populations with recurrent excitation and mutual inhibition, in the presence of independent noise sources and oscillations, see Fig 4A. This network of two competing neuronal populations may be regarded as the substrate of a number of cognitive tasks, such as perceptual bistability (visual [25,26], auditory [27], or olfactory [28]), or forced two-choice decision making [29,30].

**Fig 4.**
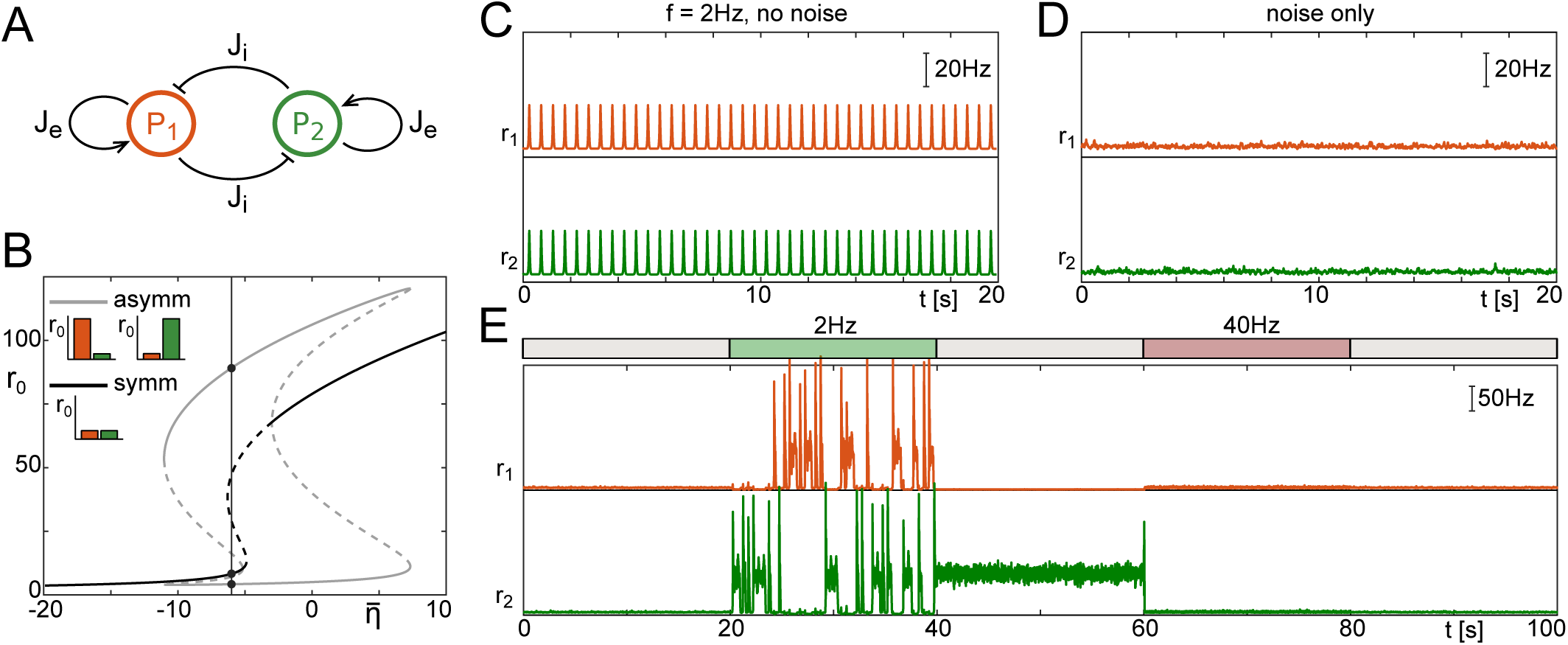
Switching in a network of two competing populations of neurons. **A** We consider two identical populations with recurrent excitatory connections and mutual inhibition. **B** Bifurcation diagram of the fixed points of the system. The system can be in a symmetric state (black) or asymmetric state (grey). We choose a point in the tri-stable regime (*η* = −6, vertical line), where either both populations are quiescent, or one population is active and the other quiescent. The insets show the stable states (two asymmetric, one symmetric). **C** Applying global forcing with slow frequency (2Hz) does not lead to the activation of either of the asymmetric patterns, due to the lack of symmetry breaking mechanisms. **D** Driving the system with independent noise sources (zero-mean Ornstein-Uhlenbeck process) with small variance *σ*^2^ does not lead to reliable switching due to long residence times. **E** Combining noise with a protocol that generates oscillations of different frequencies over different time intervals leads to the reliable (but random) activation of one of the two asymmetric patterns and switching between these at 2Hz, and the clearing of a sustained pattern at 40Hz. Parameters: 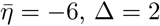 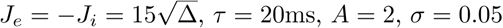.

We choose parameters such that there is one stable fixed point at which both populations are in the low firing regime, and two stable fixed points in which one population is in the low firing regime and the other is in the high firing regime. The latter two are symmetric with respect to a swap of population indices, i.e. reflection symmetric, see the bifurcation diagram in Fig 4B.

We furthermore choose an input which places the system near a sub-critical pitchfork bifurcation, but such that the symmetric, low-activity state is stable. In this state, global oscillatory forcing does not generate switching behavior at any frequency, see Fig 4C. If the two populations are driven by weak, independent noise sources, we also fail to observe any switching on relevant time scales, see Fig 4D. However, combining global oscillatory drive with weak, independent noise sources now allows for frequency-selective recall, switching, maintainance and clearance, as in the single-population network, see Fig 4D. Specifically, low frequency drive switches the network from a symmetric state to one in which one of the populations is active (2Hz stimulation in Fig 4E); continued low-frequency forcing generates ongoing stochastic switching between the two activated states. When this drive is released the currently active configuration is stabilized (between 40 and 60 seconds in Fig 4E). Finally, an intermediate range of frequencies is effective in clearing the currently held active state (40Hz stimulation in Fig 4E) and stabilizing the symmetric, low-activity state. An analysis of the bifurcation structure in this network as a function of forcing amplitude and frequency reveals that bifurcations analogous to those responsible for recall and clearance in the single-population model, i.e. Fig 3, also occur here (not shown).

#### A many-cluster network

Here we consider a network of 100 neuronal populations which interact via effective interactions which may be excitatory or inhibitory. The connectivity is chosen so that 10 distinct, random but non-overlapping activity patterns are encoded; in each pattern five neuronal populations are active, i.e. the coding sparseness is 5%. The patterns and connectivity matrix are shown in Fig 5A and B respectively. Simulations again reveal a frequency-selective response of the network similar to the two-population model. Namely, low frequency inputs in the presence of weak noise switch on the activated state and allow for robust switching, while over a range of intermediate frequencies all activated states are cleared, see Fig 5C.

**Fig 5.**
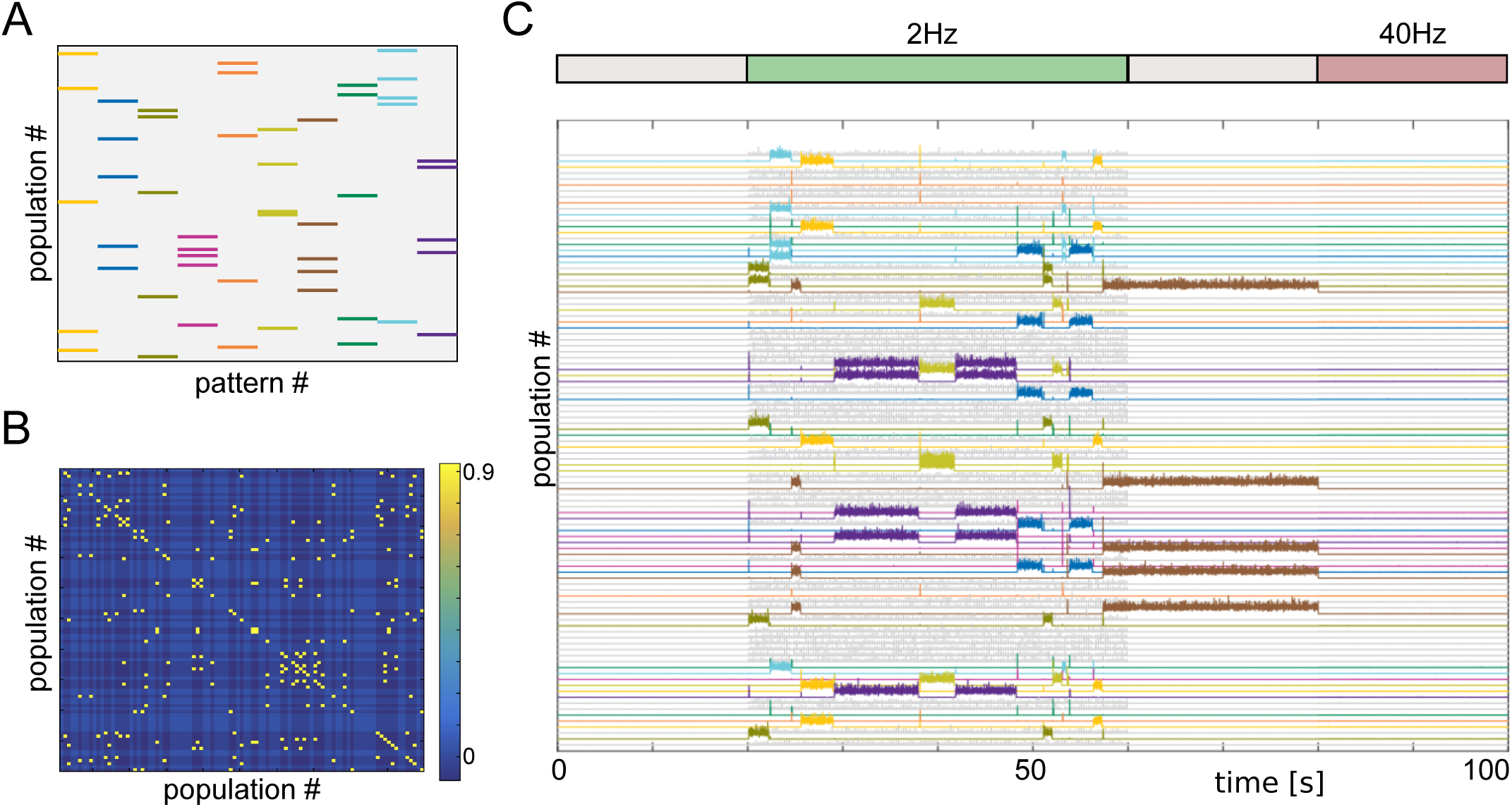
Hopfield network with random, non-overlapping patterns. **A** A network of 100 neural populations is chosen to encode ten patterns with five populations each. The patterns are non-overlapping. **B** The corresponding connectivity matrix of the network. **C** We apply an activation/deactivation protocol. The encoded patterns are randomly activated in the presence of slow oscillations (2Hz), sustained in the absence of oscillations (grey), and deactivated in the presence of fast oscillations (40Hz). All populations have independent white noise sources with variance *σ*^2^. Parameters: 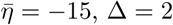 *J* = 8, *τ* = 20ms, *A* = 8, *σ* = 2.

### Generating burst-like oscillations

Thus far we have treated oscillations as an extrinsic effect, i.e. we are agnostic as to their origin. To be effective for flexible control of memory states, the oscillatory forcing we have considered here must fulfill two requirements: First, it must have a broad range of possible frequencies, and secondly, it must have a burst-like shape. Here we show that a simple circuit comprised of interacting excitatory and inhibitory populations can satisfy both these requirements.

Specifically, we construct a network of QIF neurons consisting of an E-I circuit which spontaneously oscillates, and drives a downstream population of E cells, which itself is bistable, see Fig 6. Using the corresponding mean-field equations for the E-I circuit, we found a broad region of oscillatory states of the E-I network as a function of the mean external drive to the E and I populations, *η*_*e*_ and *η*_*i*_ respectively, see the phase diagram Fig 6B. By adjusting the external drive to the E and I populations alone we can tune the output frequency over an order of magnitude. This allows us to selectively switch the downstream network on and off, as shown in Fig 6C.

**Fig 6.**
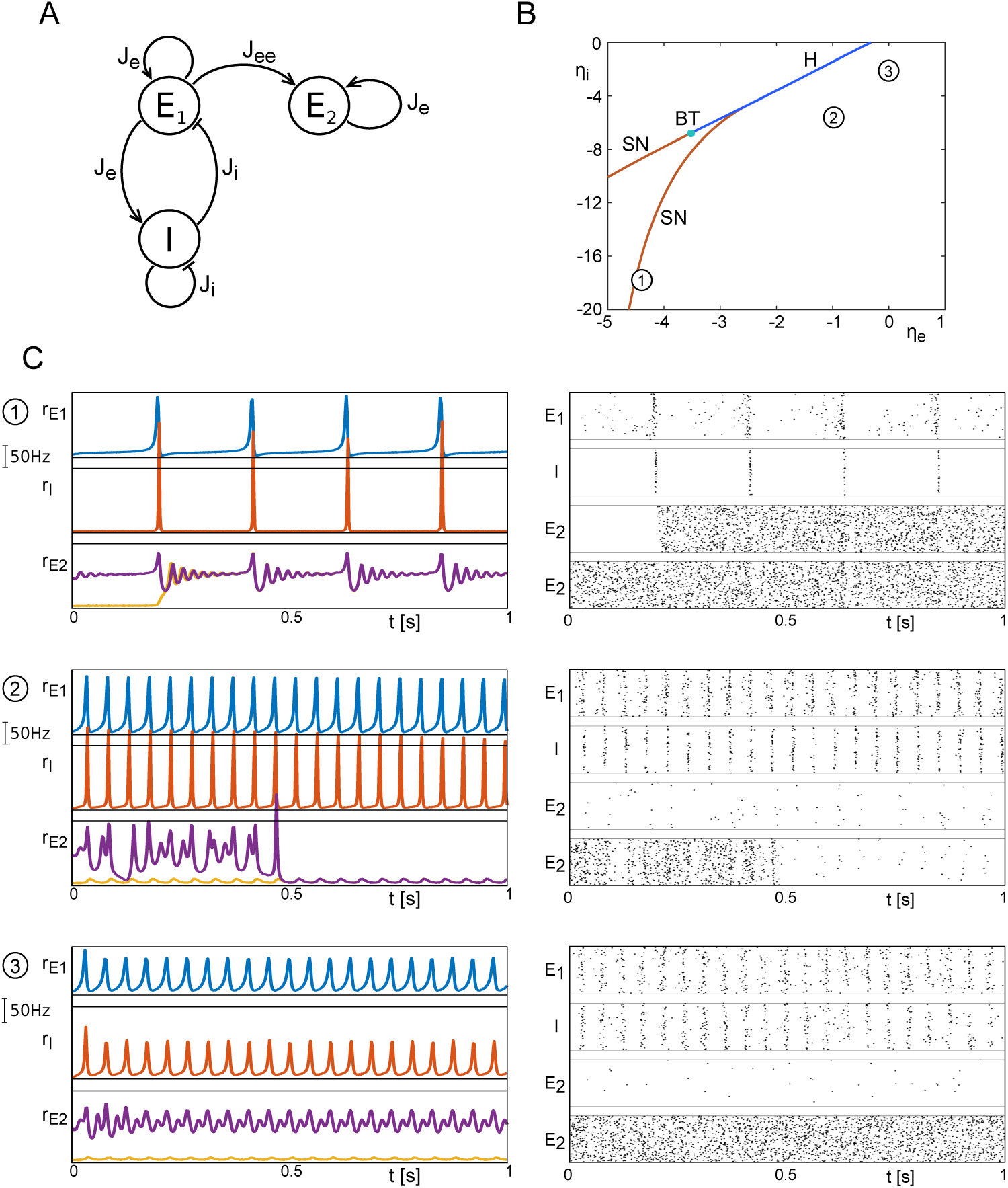
Forced switching with oscillations from an E-I network. **A** The network is built such that an excitatory population (E_1_) and an inhibitory population (I) form a circuit that can generate oscillatory output via the excitatory population (E_1_), which is fed into another excitatory populations (E_2_). The latter is in the bistable regime. **B** Bifurcation diagram of the E_1_-I-circuit in the parameters 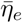 and 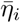The organizing bifurcations are a pair of saddle-node bifurcations (SN) of the fixed points, and a Hopf branch (H) that connects to one of the saddle-node branches via a Bogdanov-Takens codimension-two point (BT). (Limit cycles are found below the Hopf branch.) **C** Firing rates and raster plots of the population outputs as a result of the parameter tuning. Time traces of population E_2_ are portrayed for both stable initial conditions (node and focus). By choosing *η*_*e*_ and *η*_*i*_ accordingly, recall (*η*_*e*_ = *-*4.4, *η*_*i*_ = *-*18), clearance (*η*_*e*_ = *-*1, *η*_*i*_ = *-*5.5) and mai<u>nt</u>enance (*η*_*e*_=0,*η* _*i*_*= -*2) can be observed. Other parameters: 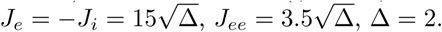. Mean current of E_2_: *η*_*ee*_ = *-*10.

### Gamma oscillations can generate memory states

Outside the region of bistability (or multistability in the case of clustered networks), neuronal networks will relax to a single stationary state in the response to a transient input. Here we show that this need not be the case if the network activity is subjected to ongoing oscillatory modulation.

As an illustration we take a single population of excitatory neurons with strong recurrent excitation, but insufficient tonic drive to place it in the region of bistability. As a result, the response of the network to a transient excitatory stimulus decays to baseline, as seen in Fig 7A (top). However, in the presence of an oscillatory input in the gamma range, which itself only very weakly modulates the network activity (Fig 7A middle), the transient input now switches the network to an activated state with prominent gamma modulation Fig 7A (bottom). Once the oscillations cease (green arrow) the activated state vanishes.

**Fig 7.**
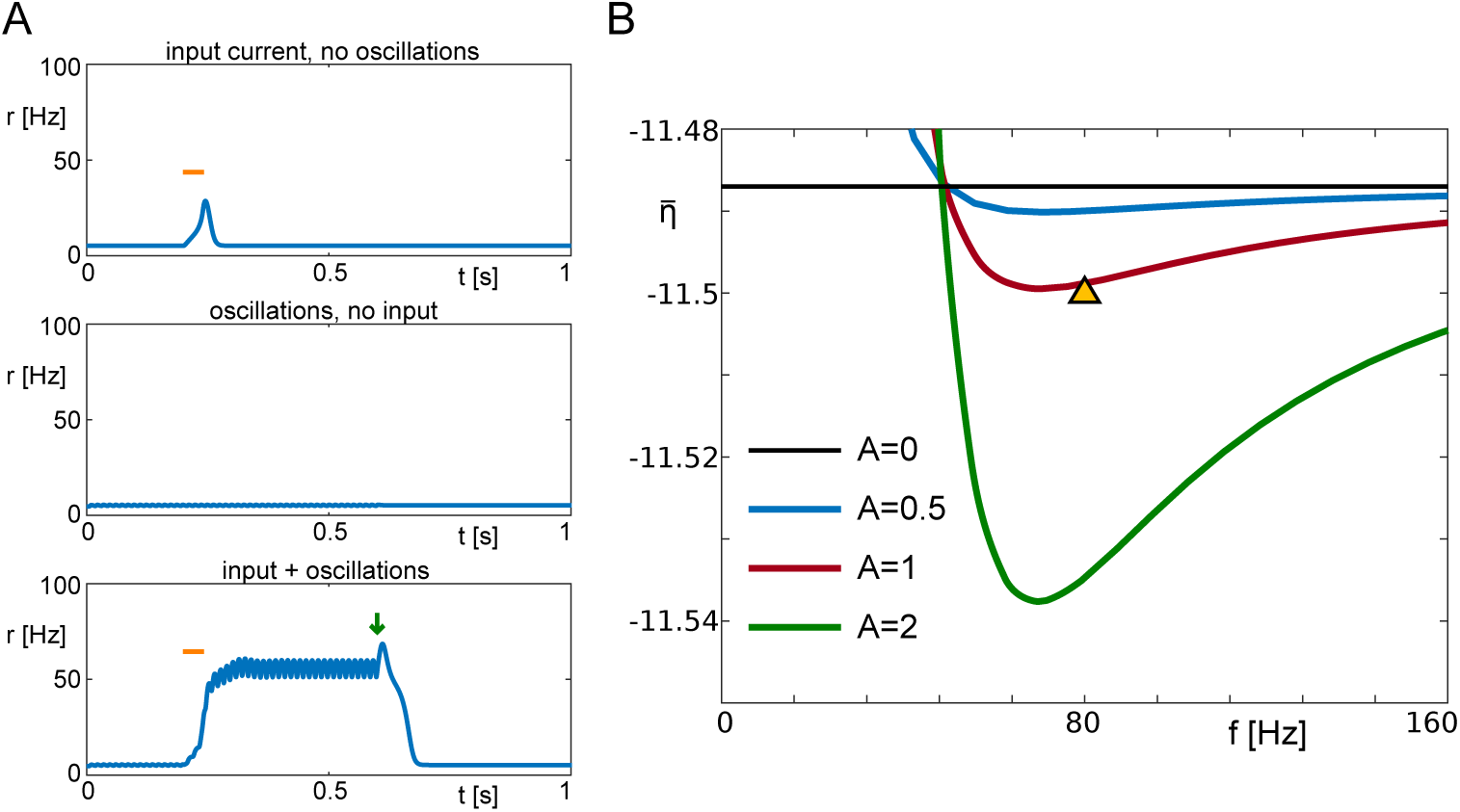
Forcing-induced bistability. **A** Here we illustrate the interplay between high frequency forcing and a transient stimulus. In the absence of oscillations, a 40ms long stimulus with amplitude 6.8 (bar) does not produce sustained activity in the model system, neither do oscillations on their own. However, the combination of oscillations with the transient stimulus leads to sustained high activity, until the oscillations are turned off (arrow). **B** Loci of the saddle-node bifurcation representing the lower limit of the bistable area, as functions of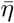and the frequency of the forcing, for different amplitudes of forcing. The choice of parameters in **A** is indicated by a triangle. Parameters: 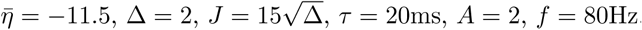.

This phenomenon depends crucially on the presence of the spike-synchrony mechanism underlying the damped oscillatory response of the high-activity focus discussed earlier. Specifically, for the parameter values used in Fig 7 the only fixed-point solution which exists is the low-activity node. Nonetheless, oscillatory forcing at sufficiently high firing rates can still recruit and resonate with the damped oscillatory interaction between the mean firing rate and mean membrane potential in the network. The resulting resonant frequency can no longer be associated with the linear response of the focus as it is a fully nonlinear network property.

The phase diagram Fig 7B shows the regions of bistability given an oscillatory forcing, for different forcing amplitudes. For zero amplitude the curve corresponds with the saddle-node (SN) bifurcation of the unforced system (horizontal black line). Note that only sufficiently high frequencies allow for bistability given tonic inputs which place the network below the SN. Furthermore, there is a clear resonance in the range of 60 − 90Hz for these parameter values. As the forcing frequency *f*→ ∞ the curves converge to the SN line of the unforced system. This is because the forcing we use has zero-mean and hence, given the low-pass filter property of neuronal networks, has no effect on the network dynamics at high frequencies.

## Discussion

In this article we study the role of oscillations in switching or maintaining specific brain states. Specifically, we identified distinct frequency bands: delta, beta, and gamma with specific functional roles. This finding is especially intriguing given that the networks we study are relatively simple. Connectivity is all-to-all and neurons are exclusively excitatory. For the multi-population networks, interactions between populations are assumed to be mediated by fast inhibition, leading to a winner-take-all behavior. Furthermore, synaptic transmission is considered to be instantaneous, with the only relevant time scale being the membrane time constant (*τ* = 20ms). The susceptibility of the networks to forcing of distinct frequencies therefore does not depend on the presence of multiple time scales associated with instrinsic currents, synaptic kinetics or sub-classes of inhibitory cells. Rather, the key dynamic factors are: bistability or multistability due to recurrent excitatory reverberation, and transient spike synchrony in response to external drive. Given this, we expect to see the same phenomenology in more biophysically realistic networks as long as there is bistability and external noise sources are not too strong.

In the region of bistability, low frequencies are effective in pushing the network into a high-activity state; for not too large amplitudes the network remains in the activated state on the downsweep of the input. The cut-off frequency for this “recall” signal is determined by the escape time of the network from the vicinity of the saddle-node bifurcation in the low-activity state, and here is a few Hertz, see Fig 2A. In multi-stable networks, this same mechanism allows for robust switching between distinct memory states. On the other hand, frequencies in the beta range are effective in switching off the high-activity state by resonantly driving bouts of spike synchrony. The precise frequency range depends on network parameters, see S4 Fig. In both cases the relevant frequency ranges scale with the membrane time constant of the neurons. Therefore, e.g. choosing a time constant *τ* = 10ms will simply stretch the x-axis of the phase diagram in Fig 2A by a factor of two. Finally, we showed that forcing in the gamma range can allow for robust working memory states which otherwise do not exist, i.e. the system sits outside the region of bistability with oscillatory forcing. This mechanism once again depends on resonantly recruiting spike synchrony.

We find that non-sinusoidal, burst-like drive is most effective in switching the network state, see Fig 3A and B. In fact, this is precisely the type of oscillation which readily emerges in a simple E-I network. Furthermore, the oscillation frequency can be modulated over a wide range through changes in the tonic drive to the E-I circuit alone, see Fig 6. This means that the state of downstream memory networks can be flexibly controlled via an E-I circuit through global changes in excitability alone.

We have proposed an E-I-circuit as a possible neuronal source of the oscillations investigated here. However, oscillations might have non-neuronal origin, such as external electric or magnetic fields during neuromodulation. Non-invasive methods include repetitive transcranial magnetic stimulation [31] and transcranial alternating current stimulation [32] which apply transient oscillatory signals. Despite the fact that they affect large parts of the brain, and hence are a challenge to model mathematically, we are confident that the methods developed here can help gain a deeper understanding of the mechanisms involved, and the psychological and behavioral effects of neuromodulation. In addition, the non-sinusoidal forcing that we introduce here might also serve to approximate the pulse-like stimulation used in deep-brain stimulation to treat Parkinson’s disease [33] and (pharmacologically) treatment-resistant depression [34].

The model we use here describes networks of spiking neurons with instantaneous synapses, i.e. the synaptic dynamics is considered fast in comparison to the membrane time scale. Future studies could incorporate synaptic dynamics with appropriate time scales for excitatory and inhibitory transmission, which can be influenced by drugs, or (pathological) changes in neurotransmitters. If synaptic time constants are in the range of the membrane time constant, we expect that the same dynamic phenomena observed here still occur. If synaptic time constants are large, however, spike synchronization may no longer occur [23]. The framework developed here may therefore serve as a tool to study the cause of functional deficiencies in synapse-related conditions, so-called ‘synaptopathies’ [35,36]. In addition, one may envisage more detailed cortical network architectures involving multiple neuronal populations [37–39]. In this article, we briefly investigated the role of an E-I-circuit to generate oscillations which control the state of another neural population. We are therefore optimistic that using realistic circuit models in combination with the mean field description employed here will yield new and interesting results.

## Materials and Methods

### Mathematical Model

An important mathematical tool in understanding macroscopic neuronal dynamics, away from the description of spiking neurons, is provided by neural mass and neural field models. Classical models include the Wilson-Cowan model [40, 41] or the Amari model [42, 43]. However, such macroscopic models of brain activity often pose a stark simplification of the actual dynamics, and often miss important features from the spiking dynamics, such as spike synchronization as shown in this article. Therefore, there have been recent advances in linking the microscopic and macroscopic dynamics of networks of spiking neurons [15, 44–53].

We consider a neural mass model that was recently derived from networks of all-to-all coupled quadratic integrate-and-fire neurons in the thermodynamic limit [15], see Eqs (1). To simplify the mathematical treatment, we divide *t* by *τ* which represents the case of time being measured in units of *τ*, thus eliminating *τ* from the equations:

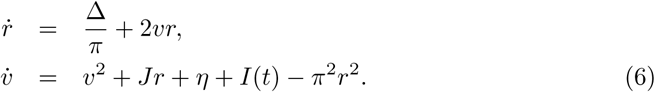

Here, *r* represents the ensemble average of the firing rate of neurons, and *v* represents the ensemble average of the membrane potential. The parameters *η* and Δ represent the mean and variance of the Lorentzian distribution of time-invariant input currents into the neuronal ensemble, and *J* is the coupling constant between neurons. Time-varying external inputs are given by *I*(*t*). The original model (1) can then be recovered by *t→ τt, r →r/τ*. As we set *τ* = 20ms, *r* = 1 here corresponds to a firing rate of *r* = 50Hz in the full model.

Here, we consider *I*(*t*) to be *T*-periodic, i.e. *I*(*t* + *T*) = *I*(*t*). We distinguish between two types of input: sinusoidal input,

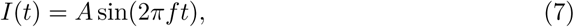

and non-sinusoidal input,

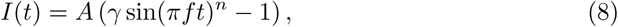

where we take *n* = 20 for the simulations presented in this paper. The parameter *A* represents the amplitude of the forcing. The constant *γ* is chosen such that 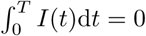. We choose this type of zero-mean forcing to avoid any changes in network excitability which a tonic DC-offset might cause. In other words, the input models a reorganization of afferent spikes into periodic volleys without adding any additional spikes. In the non-sinusoidal case the spikes are more synchronized than in the sinusoidal case.

We compare the full model equations with its equivalent heuristic firing rate equation, which preserves the fixed point structure but reduces the dynamical behavior. This is done by considering stationary solutions given by

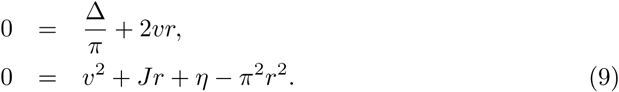

Solving these equations for *r* is equivalent to solving Eq 2. Thus, the reduced heuristic firing rate equations can be expressed by

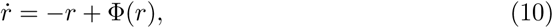

where the f-I function Φ(*r*) is given by Eq 3. Straightforward stability analysis gives the stability condition Φ *′*(*r*) *<* 1.

### Linear Response

Ignoring transient dynamics, the response of the model equations to the external input *I*(*t*) is *T*-periodic as well, at least in the limit of small amplitudes *A* (an exception are period-doubled solutions, which are a nonlinear phenomenon only relevant at larger *A*). In this case the corresponding Fourier spectra of the firing rate *r*(*t*) and of the membrane potential *v*(*t*) are discrete:

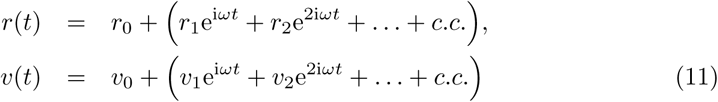

For brevity of exposition we use here the angular frequency *ω* = 2*πf* instead of the ordinary frequency *f*. This approach describes the projection of solutions of *r* and *v* from a continuous space ℝ onto a discrete function space *V*, with orthogonal basis functions e^i*nωt*^,*n*∈ℤ. The same Fourier decomposition applies to the input current *I*(*t*):

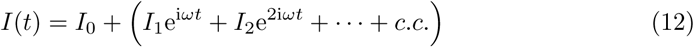

To determine the linear response of the model equations, we first carry out Fourier decomposition of the linearized system:

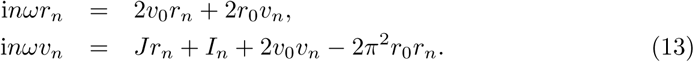

Solving this set of linear equations, we obtain

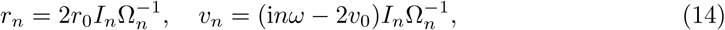

with

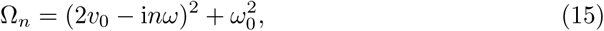

where *ω*_0_ is the (angular) resonant frequency:

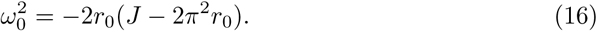

The resonant frequency is state-dependent and changes with model parameters. Reintroducing the time scale *τ*, perturbations of the upper branch solution resonate at a frequency

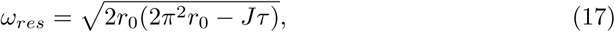

where *r*_0_ is the value of the steady-state firing rate. This is true as long as the argument of the square root is positive. Therefore as the firing rate decreases along the upper branch, for decreasing external input, the frequency decreases to zero at which point the focus becomes a node. This point occurs before the saddle-node is reached unless Δ = 0 in which case it exactly coincides with the saddle-node.

The time-dependent linear response of the firing rate and the membrane potential is now given by

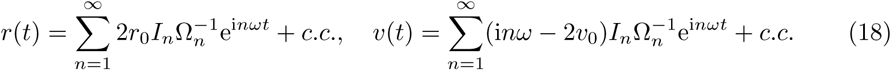

From this, we can derive the amplitude of the linear response of the firing rate,

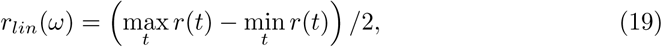

and analogously of the membrane potential. Alternatively, one can derive the time-averaged linear response (“power”) of the system:

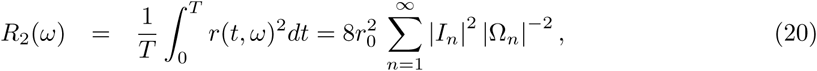

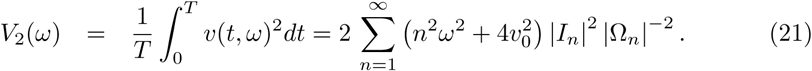

Here we have made use of the orthogonality of the basis functions, and the fact that *T* = 2*π/ω*.

### Numerical Continuation

In order to exhaustively and accurately trace the bifurcations that occur in the model equations, we make use of AUTO 07p [54]. Since this software is designed to deal with autonomous systems, we recast the (non-autonomous) model equations (1) as a set of autonomous equations:

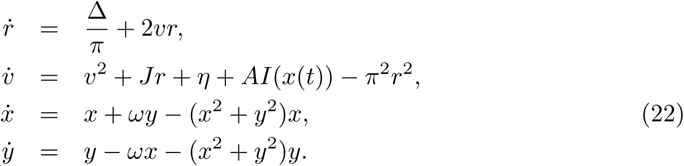

The last two equations create the periodic stimulus *x*(*t*) = sin(*ωt*) in the model equations. We distinguish the sinusoidal case,

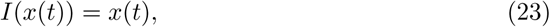

and the non-sinusoidal case

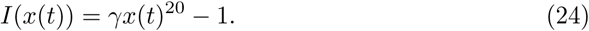

Continuation of the forced system is performed by starting from a known fixed point (*r*_0_, *v*_0_) at *A* = 0, and continuing solutions by increasing *A* up to the desired value. We use the *L*_2_-norm as a scalar measure to represent periodic solutions:

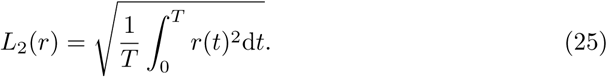

Where we perform this one-parameter continuation, we represent solution branches by plotting the *L*_2_-norm against the parameter that is being varied. Where we perform two-parameter continuation, we plot the loci of bifurcations against the two parameters being varied.

### Memory Networks

A natural extension of the single-population model is to consider a network of neural masses:

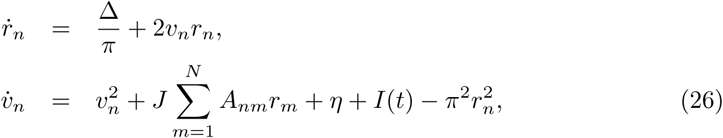

where the adjacency matrix *A* determines the connectivity structure between neural masses. In this paper we consider two scenarios, the first of which is two neural populations with recurrent excitation and mutual inhibition. The adjacency matrix of such a network is given by

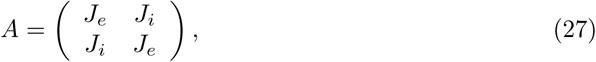

where *J*_*i*_ *<* 0 *< J*_*e*_.

In the second scenario, we examine the dynamics within a Hopfield network. Rather than creating the network through learning algorithms, we build the network as follows. First, we choose the patterns that the network should encode and write them into an arrayÛ. Each column of this array represents one pattern, where we put 1 for populations that are active in this pattern, and 0 otherwise. As a result, the array Ûhas the size *N* × *N*_*pat*_, where *N*_*pat*_ is the number of patterns encoded, and *N* is the network size. Each pattern consists of *N*_*p*_ active populations, and patterns are non-overlapping. The adjacency matrix of a network that encodes these patterns then reads

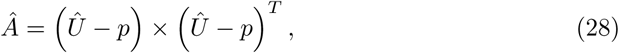

with *p* = *N*_*p*_*/N*.

### E-I circuit generating oscillations

To create a network that generates oscillations, we consider a network of an excitatory population interacting with an inhibitory one:

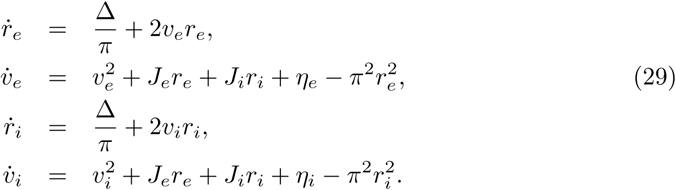

For simplicity, we choose *J*_*e*_ = *-J*_*i*_ = *J*. The two populations differ in terms of the means of their tonic input currents, *η*_*e*_ and *η*_*i*_. We vary these two parameters to identify the regime where stable oscillations exists, and to change the frequency of these oscillations.

## Supporting Information

### S1 Text. Mechanisms underlying switching

Here we illustrate in greater detail the mechanisms underlying the “switching on” of activated network states (or simple switching between attractors in the case of a multi-stable network) at low frequencies, and the “switching off” of activated states at frequencies in the beta range.

The effect of low frequencies can be understood by considering the quasi-stationary response, namely how changes in the external drive alter the steady-state network solution. S1 FigA-C show the steady-state bifurcation diagram for a single excitatory population of QIF neurons as a function of the mean external drive *η*.

**S1 Fig.**
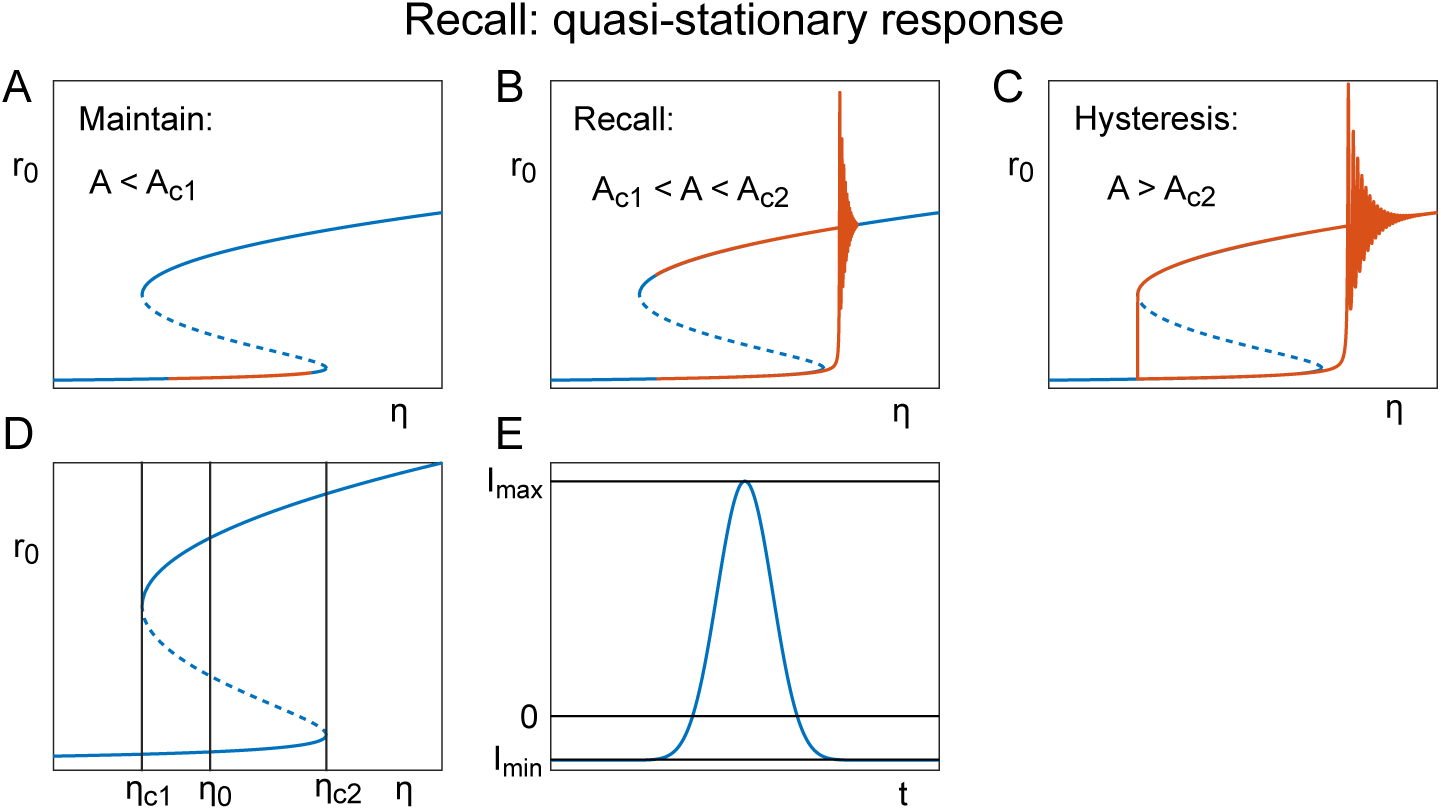
Mechanisms of switching: Quasi-stationary response. **A** At amplitudes below the critical range no switching occurs (*A* = 0.7). **B** Amplitude values within the critical range lead to switching (*A* = 1). **C** At amplitudes above the critical range the system undergoes periodic hysteretic switching (*A* = 1.3). **D** Bifurcation diagram of stationary states with critical values for saddle-node bifurcations (*η*_*c*1_, *η*_*c*2_) and the choice of model parameter (*η*_0_). **E** Normalized non-sinusoidal forcing over one period (*A* = 1), with minimum and maximum values indicated. Parameters: 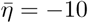 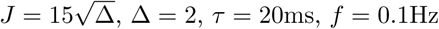.

On top of these bifurcation diagrams we plot the firing rate of the forced system against the *x*-axis, which is 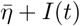 as the forcing can be understood to be a time-varying mean input current into the system. This is to illustrate that at low frequencies the system remains close to the fixed points (except when it switches between them), hence the term ‘quasi-stationary’. Given a mean input which places the system within the region of bistability, a small-amplitude, low-frequency forcing fails to push the system past the low-activity saddle-node, see S1 FigA. In a range of forcing amplitudes the network switches to the high-activity state and remains on the upper solution branch, see S1 FigB, while for larger amplitudes the network activity becomes slaved to the forcing, S1 FigC. The range of suitable amplitudes depends on the model parameter *η*, which is situated in the bistable regime. For clarity, we denote the chosen parameter by 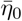. The bistable regime is delimited by two saddle-node bifurcations, that occur at 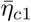 and 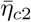, respectively. Thus we have 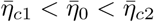The range of amplitudes also depends on the shape of the forcing representative of the case *A* = 1. In this case, the forcing is characterized by its minimum value *I*_*min*_ and its maximum value *I*_*max*_. We assume *I*_*min*_ *<* 0 *< I*_*max*_. The minimal amplitude required to push the system from the node to the saddle is then given by

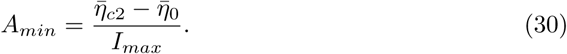

The maximum amplitude, up to which the system stays on the upper branch, is given by

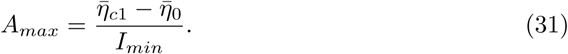

If the system parameters are such that *A*_*max*_≤*A*_*min*_, then there is no amplitude regime at which Recall occurs, and increasing the amplitude leads to a transition from Maintenance directly to hysteresis.

S2 Fig shows the details of the bifurcation structure of the periodically forced network which leads to the “switching off” or “clearance” behavior.

**S2 Fig.**
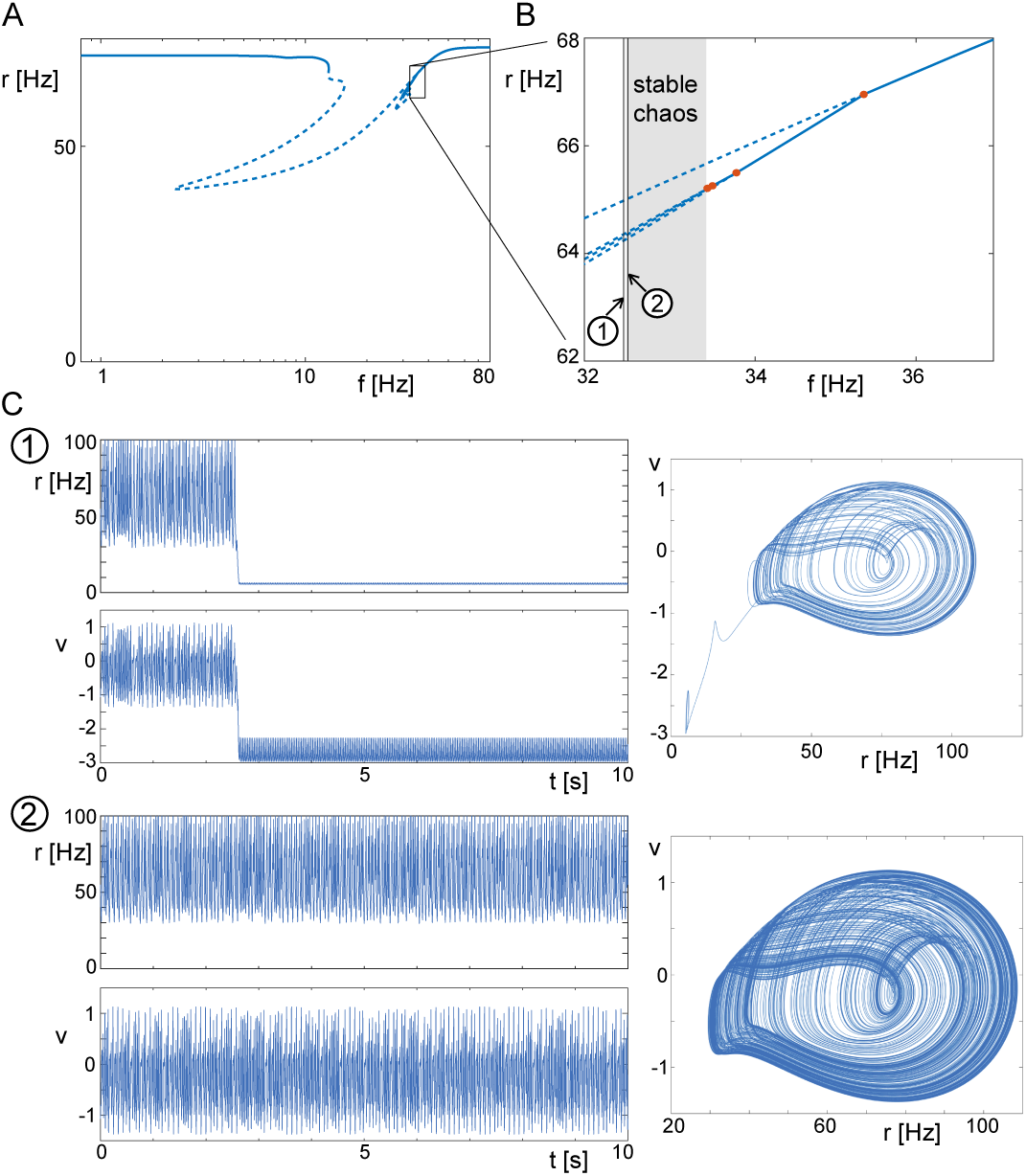
Mechanisms of switching: Nonlinear resonance. **A** Bifurcation diagram of the focus with *f* as bifurcation parameter at *A* = 1. **B** Inset of **A**, with period-doubling bifurcations (orange dots) and emerging branches of period-doubled solutions shown. The period-doubling cascade gives rise to stable chaos (grey area), which becomes unstable at lower frequencies. **C** Example time series from **B** around the area where the chaotic attractor becomes unstable.

Specifically, a series of period-doubling bifurcations, a so-called period-doubling cascade, leads to the emergence of a chaotic orbit. This orbit is initially stable, but a further decrease of the frequency leads to global instability of the chaotic orbit, and the destabilization of the high-activity limit cycle solution, see S2 FigA-B. The latter occurs just below a forcing frequency of *f* = 32.5Hz, see S2 FigB. In S2 FigC we show representative time traces for forcing frequencies of *f* = 32.45Hz and *f* = 32.5Hz. In the former case, the system leaves the forced focus in less than three seconds, whereas in the latter case the chaotic orbit persists for the whole simulation period of 10^3^ seconds. We infer from this that the critical frequency at which the chaotic orbit loses stability globally is within this frequency range.

### S2 Text. A canonical model for nonlinear resonance in the bistable regime

In the network model, the high-activity branch of solution in the bistable regime exhibits damped oscillations. Periodic external drive can resonate with these intrinsic oscillations, leading to destabilizing period-doubling bifurcations as seen in the previous section. Here we show that this mechanism is present in the simplest possible model exhibiting a saddle-node bifurcation and for which the upper branch becomes a focus beyond a critical value of the external drive:

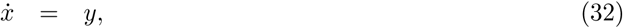

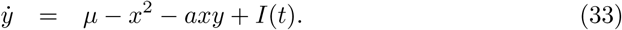

This model is a particular unfolding of the so-called Takens-Bogdanov normal form, for which there is no Hopf bifurcation, which is the relevant case for our network model. It is easily shown that a saddle-node bifurcation occurs in these equations at *μ* = 0 and that the fixed point solutions are 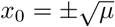 and *y*_0_ for μ > 0, see S3 FigA.

**S3 Fig.**
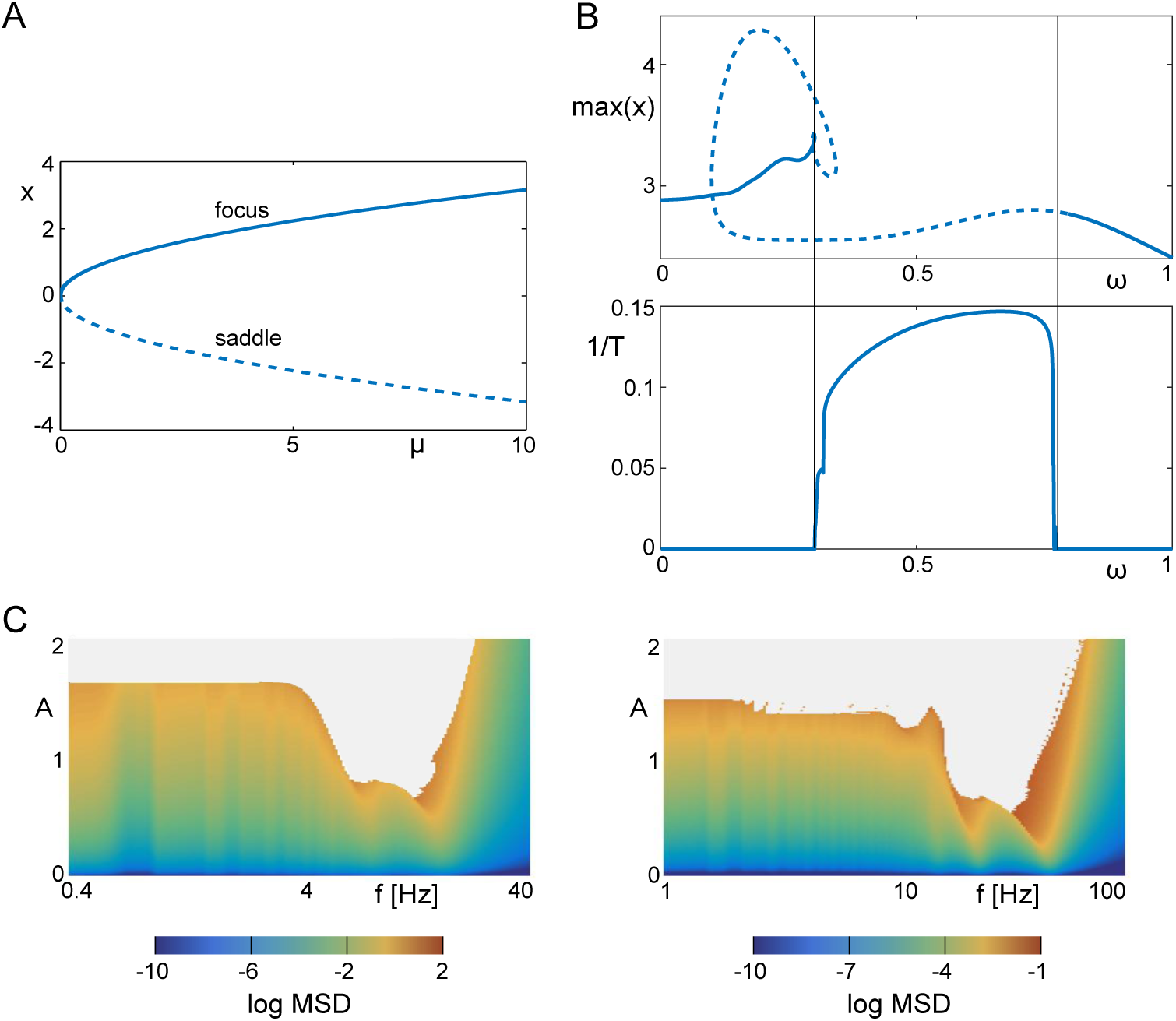
Fig The nonlinear resonance is captured in a canonical normal form for a saddle-”focus”. **A** Bifurcation diagram of fixed points of this system, giving rise to a stable focus and an unstable saddle. **B** *Top:* The bifurcation structure of solutions resulting from non-sinusoidal forcing with *f* as bifurcation parameter (*A* = 1). The area between vertical bars contains unstable period-doubled solutions (not shown), which is evidence of the existence of a chaotic attractor. *Bottom:* Inverse of the time *T* that *x* needs to reach an absolute value of 10^6^, which is evidence that solutions diverge due to an unstable chaotic attractor. **C** Comparison of the nonlinear response of the reduced system (left) with the full system (right). Parameters: *μ* = 2, *a* = 0.4.

Furthermore, the solution 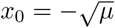 is a saddle, and 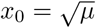 is a stable focus for which the frequency goes to zero smoothly as *μ →*0. S3 FigB shows that in the forced system there is a range of frequencies for which there is no stable solution; in the normal form equation the solution diverges while in the network model the system settles to a limit cycle solution in the vicinity of the low-activity state. The instability is due to a series of period-doubling bifurcations as in the full system. Furthermore, comparison of the phase diagram of the normal form equation with that of the full system shows they are qualitative similar, S3 FigC. This indicates that the nonlinear resonance seen in the network of QIF neurons is a generic feature of any system with a stable focus in the vicinity of a saddle-node bifurcation.

### S3 Text. How the resonant frequency changes with network parameters

S4 Fig shows how the linear resonant frequency of the stable focus in the bistable regime of a network of excitatory QIF neurons varies as a function of the mean external input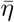and the strength of synaptic coupling *J*. Recall is not possible to the left of the red curve given the nonlinear forcing used here. This line is determined by setting *A*_*min*_ = *A*_*max*_, see Eq (30) and Eq (31).

**S4 Fig.**
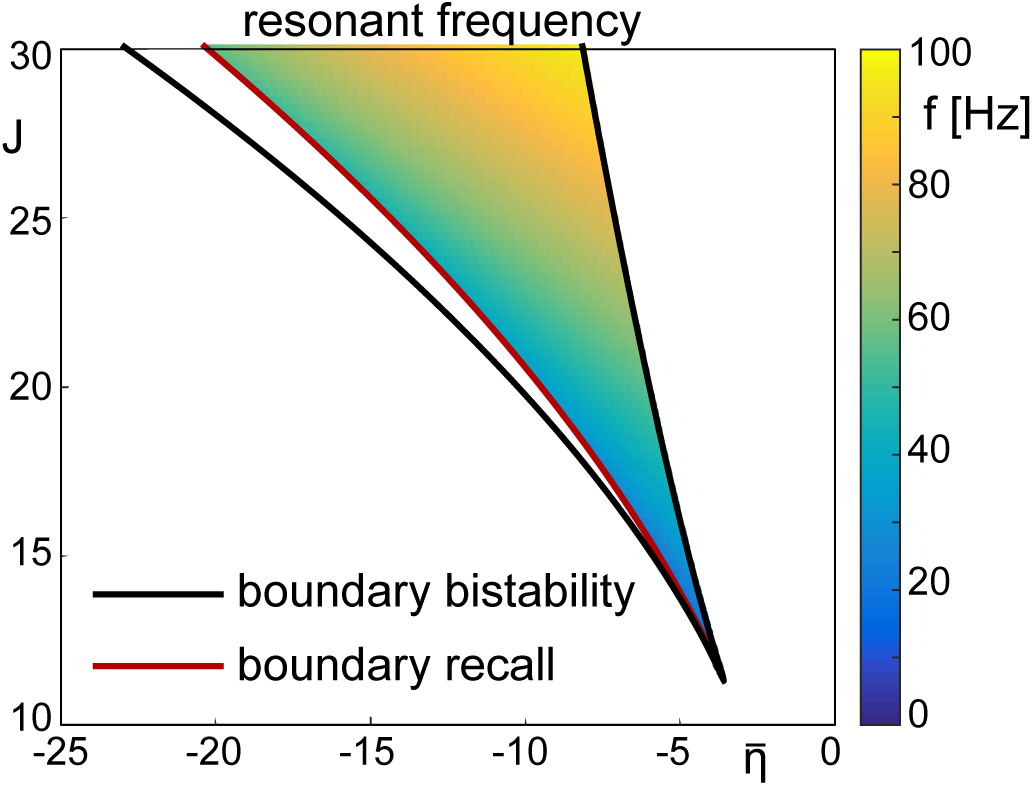
Change of resonant frequency with model parameters. We compute the (linear) resonant frequency of the saddle for parameter values in the bistable regime (delimited by black lines), specifically where Recall occurs using non-sinusoidal forcing (delimited by red line). As Clearance is caused by nonlinear resonance of the focus, the corresponding frequency band is found near the linear resonant frequency. At fixed values of *J* the resonant frequency varies approximately by a factor of two across the range of values of 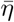Other parameters: Δ = 2, *τ* = 20ms.

